# Impairing flow-mediated endothelial remodeling reduces extravasation of tumor cells

**DOI:** 10.1101/2020.07.27.219568

**Authors:** Gautier Follain, Naël Osmani, Luc Mercier, Maria Jesus Garcia-Leon, Ignacio Busnelli, Angélique Pichot, Nicodème Paul, Raphaël Carapito, Siamak Bahram, Olivier Lefebvre, Jacky G. Goetz

## Abstract

Tumor progression and metastatic dissemination are driven by cell-intrinsic and biomechanical cues that favor the growth of life-threatening secondary tumors. We recently identified prometastatic vascular regions with blood flow profiles that are permissive for the arrest of circulating tumor cells. We have further established that such flow profiles also control endothelial remodeling, which favors extravasation of arrested CTCs. Yet, how shear forces control endothelial remodeling is unknown. In the present work, we aimed at dissecting the cellular and molecular mechanisms driving blood flow-dependent endothelial remodeling. Transcriptomic analysis revealed that blood flow modulated several signaling pathways in endothelial cells. More specifically, we observed that VEGFR signaling was significantly enhanced. Using a combination of in vitro microfluidics and intravital imaging in zebrafish embryos, we now demonstrate that the early flow-driven endothelial response can be prevented with sunitinib, a pan-inhibitor of VEGFR signaling. Embryos treated with sunitinib displayed reduced endothelial remodeling and subsequent metastatic extravasation. These results confirm the importance of endothelial remodeling as a driving force of CTC extravasation and metastatic dissemination. Furthermore, the present work suggests that therapies targeting endothelial remodeling might be a relevant clinical strategy in order to impede metastatic progression.

**Highlights:**

- Flow profiles that are permissive for metastasis stimulate the VEGFR pathway
- Flow-dependent VEGFR signaling favors extravasation of CTCs through endothelial remodeling
- Inhibition of VEGFR signaling suppresses early flow-driven endothelial response

## Introduction

Metastatic colonization occurring during advanced tumorigenic progression is the main reason for cancer-related death (Massagué and Obenauf, 2016). It is currently well admitted that their development relies on biological and biophysical cues that influence their location and development (Follain et al., 2019). Indeed, adhesion molecule repertoire (Reymond et al., 2013; Osmani et al., 2019), vascular architecture (Chambers et al., 2002; Kienast et al., 2010; Headley et al., 2016) and hemodynamics (Wirtz et al., 2011; Follain et al., 2018a, 2020) impact the early steps of tumor cells dissemination. Several biochemical and physical parameters found in colonized organs such as growth factors, chemokines, stromal cell composition (Paget, 1889; Obenauf and Massagué, 2015; DeNardo et al., 2010; Nagarsheth et al., 2017) and matrix stiffness (Paszek et al., 2005; Levental et al., 2009; Mouw et al., 2014) will permit the growth of secondary tumors. In order to reach distant organs, tumor cells need to go through several rate-limiting steps including intravasation, blood-borne transport and extravasation (Valastyan and Weinberg, 2011). Thus, a comprehensive understanding of these steps is necessary in order to design relevant pharmacological therapeutic strategies. Recently, using zebrafish embryo and mouse experimental metastasis models, we have demonstrated that blood flow-induced endothelial remodeling acts as a major driving force to cell extravasation and subsequent micro-metastasis formation (Follain et al., 2018a). Specifically, we have shown that the endothelium wall actively remodels around arrested tumor cells in order to maintain vessel perfusion and thus actively promotes tumor cell extravasation. Such phenomenon relies on the ability of endothelial cells to protrude apically, migrate intravascularly and enwrap arrested CTCs, excluding them from the inner vasculature. It was suggested that MMP enzymatic activity is required to resolve blood clot and maintain vessel perfusion through endothelial remodeling in mouse brain (Lam et al., 2010). However, the molecular mechanisms driving endothelial remodeling downstream of blood flow biomechanical cues remain elusive. In the following report, we aimed at identifying the molecular programs that are transcriptionally activated downstream of blood flow, at values that are permissive to metastatic extravasation. Among others, activation of VEGFR signaling emerged as an attractive target and a potential major endothelial remodeling signature. Using sunitinib, a wide spectrum tyrosine-kinase and VEGFR inhibitor approved for clinical use for treating renal cell carcinoma (Carmeliet and Jain, 2011), we further demonstrate that VEGFR inhibition impairs extravasation of circulating tumor cells by blocking the endothelial remodeling. Thus, our findings confirm that endothelial remodeling is an essential step for metastatic extravasation and can be molecularly controlled using existing pharmacological tools (Meadows and Hurwitz, 2012).

## Results

### Flow upregulates VEGF signaling pathway

In order to identify signaling pathways that drive endothelial remodeling in dependence of permissive flow forces, we applied RNA sequencing on cultured primary endothelial cells (HUVEC) to identify transcriptional programs activated by previously identified flow profiles. We used previously described methods based on Human Umbilical Venous Endothelial Cells (HUVEC) culture in microfluidic channels (Follain et al., 2018a). The HUVEC monolayer were cultured for 16 hours in static conditions or with medium perfusion a flow rate of 400 μm/sec (Fig 1A). Such flow velocity was selected for two reasons. Such flow regime favors extravasation of tumor cells in zebrafish embryos and it matches the flow rate measured in capillary like vessels that we and others reported (Ivanov et al., 1985; Anton et al., 2013; Hill et al., 2015; Sugden et al., 2017; Follain et al., 2018a). Total RNAs were then extracted, quantitative RNA sequencing was carried out and flow-responsive genes were clustered and functionally annotated using Gene Ontology (GO) databases. Strikingly, endothelial cells switched their transcriptional programs from cell division and mitosis to cell migration/angiogenesis when exposed to flow (Fig 1B and Fig S1A), as previously suggested (Fang et al., 2017). Interestingly, a significant fraction of genes upregulated in response to flow is involved in vascular development, angiogenesis, and response to oxidative stress (Fig 1B and Fig S1A). This is in accordance with previously described endothelial transcriptional response to shear stress (Simons et al., 2016). Shear stress is indeed an essential biomechanical cue that controls sprouting and migration of endothelial cells sprouting as vessel remodeling during angiogenesis (Galie et al., 2014; Baeyens et al., 2015; Baeyens and Schwartz, 2016). We thus hypothesized that angiogenesis related genes might be elemental in driving endothelial remodeling during CTC extravasation. We thus focused on the GO class “Angiogenesis” (Fig 1C) and applied a stringent threshold to our list of significantly impacted gene (with fold changes > 1.5 and p-value < 0,05). Among others, we identified FLT1 (VEGFR1) and KDR (VEGFR2), two VEGF receptors (VEGFR) that were strongly upregulated in endothelial cells subjected to our reference flow velocity (400 μm/sec). Using RT-qPCR, we validated a two-fold increase in gene expression of FLT1 and KDR upon flow stimulation (Fig S1B). Using similar conditions, we assessed the protein levels of all three VEGFRs (FLT1, KDR and FLT4 (VEGFR3)) using western-blot and immunofluorescence (Fig 1D-F). Expression levels of KDR are significantly and consistently increased in response to flow. Expression levels of FLT1 and FLT4 also increase in response to flow, yet they display a heterogeneous response (Fig 1D-F). Altogether, these data suggest that permissive flow profiles for endothelial remodeling and subsequent extravasation of CTCs drive expression of VEGFRs, both at the RNA and protein levels. This prompted us to test the role of VEGFR pathways on the flow-dependent endothelial remodeling leading to extravasation of tumor cells.

**Fig. 1.**
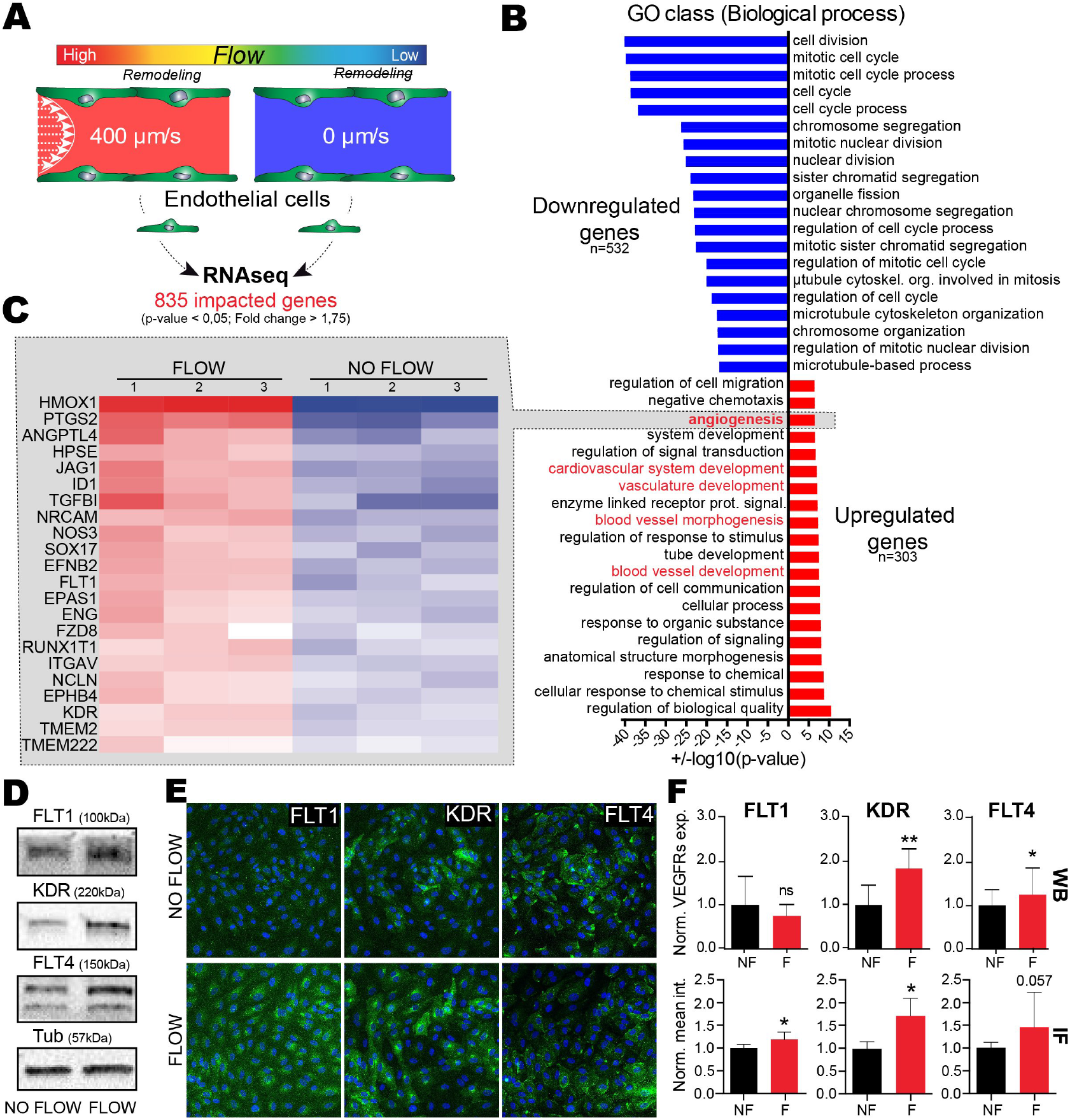
Flow favors expression of gene program driving endothelial remodeling, including VEGFRs. A - Experimental setup: microfluidic experiment for RNA sequencing. B - Results from global Gene Ontology analysis on ‘biological process’, showing most significantly impacted GO class. C – Fold changes heatmap, based on GO class: Angiogenesis, showing significantly upregulated genes in flow condition compared to no flow. D – Western blot quantification validating of some of VEGFR RNAseq data. E – Representative image of flow vs no flow immunofluorescence labelling against FLT1, KDR and FLT4 (green) with DAPI (blue). F – Quantification associated with D and E, showing increased expression and signal intensity of VEGFRs. WB: N=5, Mann Whitney test, IF: N=4 with min 5 slices/exp, Mann Whitney test.

### Inhibition of the VEGF pathway in vitro suppresses endothelial remodeling

Based on our previous results, we hypothesized that VEGFR signaling could trigger flow-dependent endothelial remodeling, and subsequent metastasis, and decided to apply to pharmacologically impair it. Here, in a proof-of-concept approach, we decided to employ pharmacological inhibition using a clinically approved drug, sunitinib (Sutent® or SU11248), for its known anti-angiogenic properties through VEGFRs, among others. Our goal, at this stage, was to demonstrate that molecular inhibition of flow-mediated endothelial remodeling is a possible avenue for impairing metastatic extravasation. We first confirmed that VEGFR2, a VEGFR isoform we found consistently overexpressed at the RNA and protein level, is recruited at the plasma membrane of endothelial, cells undergoing remodeling of CTC in vitro (Fig 2A). Using similar microfluidics where endothelial cells were subjected to VEGFR inhibition using sunitinib prior to CTC perfusion, we next assessed whether such treatment would impact endothelial remodeling and CTC transmigration (Fig 2B). The microfluidic channels were perfused with tumor cells (CTCs) that adhered to endothelial cells and further flow-stimulated for 16 hours in the presence of vehicle or sunitinib. As previously described (Follain et al., 2018a), flow stimulation favored transmigration through the endothelial cells monolayer of 95 per cent of the cancer cells. Among these, the vast majority (80 per cent) of the cells that had crossed the endothelial wall did so through endothelial remodeling (Fig 2C-D). VEGFR inhibition through Sunitinib treatment significantly decreased the number of tumor cells that could cross the endothelial barrier and suppressed the endothelial remodeling activity (Fig 2C-D). In conclusion, flow stimulates VEGFR-dependent endothelial remodeling and subsequent metastatic extravasation in microfluidic artificial environments in vitro.

**Fig. 2.**
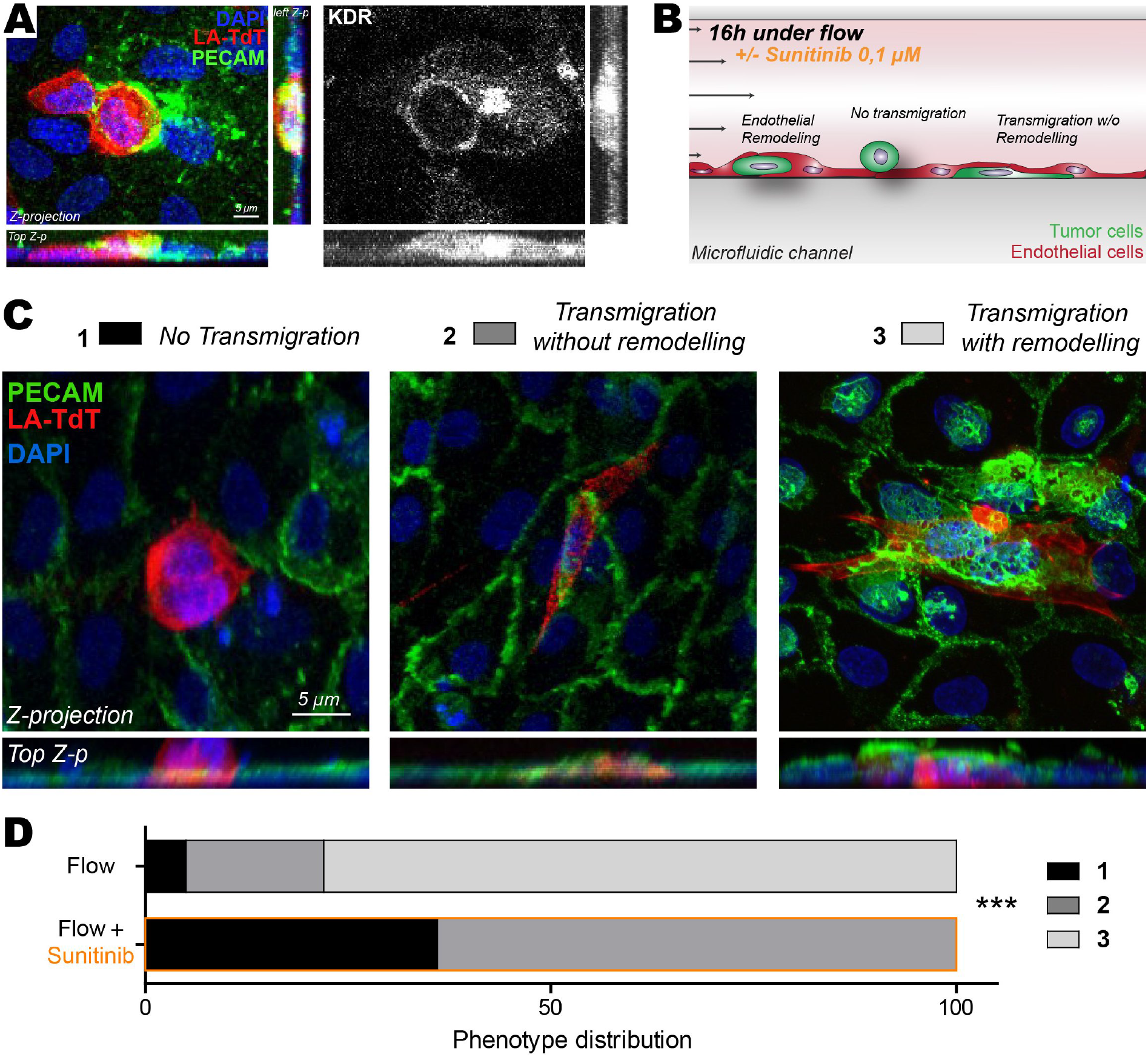
Inhibition of VEGFRs with sunitinib impairs endothelial remodeling in vitro. A – Immuno-labelling pictures showing the presence of KDR (white) at the site of endothelial remodeling (Assessed with PECAM enrichment (green) around tumor cell (red)). Z-projections and two z-projection from side and top view are shown. B - Experimental setup: microfluidic channel for endothelial remodeling assay. C – Representative images of all steps: 1-No transmigration, 2-Transmigration without remodeling evidence, 3-Transmigration with remodeling. Quantification of the normalized number of transmigrating cells and remodeling activity. D – Phenotypic distribution of CTCs attached to endothelial layers exposed to flow and treated with vehicle or sunitinib. N: flow/vehicle = 88, flow/sunitinib = 150, Chi2 test.

### Inhibition of the VEGFR pathway inhibition blocks early endothelial remodeling in zebrafish embryos

We next wondered whether such mechanism would occur in realistic hemodynamic situations. Zebrafish is emerging as an easy-to-use experimental metastasis model which is fully compatible with intravital imaging and chemical compound screening which can be directly added into the water (Cagan et al., 2019; Osmani and Goetz, 2019). We thus set out to test the involvement of VEGFR signaling in endothelial remodeling in vivo, using intravital imaging of intravascularly arrested tumor cells in zebrafish embryos. To rule out side-effects, we first controlled whether a short (3h) sunitinib treatment (5μM) would impair the vascular system and hemodynamics of Tg(fli1a:EGFP; gata1:RFP) zebrafish embryos at 48 hours post-fertilization (hpf) expressing EGFP in the endothelium and RFP in red blood cells (Fig 3A). Pharmacological treatments of the embryos had mostly no impact on the vascular architecture of embryos, with only the most caudal inter-somitic vessels (ISVs) failing to lumenize in a few embryos (Fig 3A). Anatomical analysis and quantification of the vascular system revealed that the vascular architecture was mostly unperturbed upon sunitinib treatment (Fig 3B). Furthermore, using automated tracking analysis of the blood flow (based on dynamic imaging of RFP-positive red blood cells) that allows to extract and quantify the mean, minimum and maximum velocities as well as pulsatility of the blood flow, we could not detect any major effect of VEGFR inhibition of overall hemodynamics (Fig 3C). Altogether, this set of control experiments demonstrate that short sunitinib treatment does not impact the pre-established vascular system, allowing us to investigate whether it would perturb endothelial remodeling and subsequent metastasis. Using our intravital imaging-based experimental metastasis model in zebrafish, we next injected tumor cells directly in the circulation through the Duct of Cuvier of 48hpf embryos (Follain et al., 2018b) and quantitatively address the effect of sunitinib on extravasation in vivo (Fig 3D). We classified the behavior of injected tumor cells into three populations at different stages of this process, whose dynamics has been characterized in our previous work: “intravascular”, “pocketing” (i.e. in the process of extravasation through endothelial remodeling) and “extravasated” (Follain et al., 2018a). After only 3 hours post-injection (hpi), tumor cells are stably adhered to the endothelium, which starts to remodel in order to restore blood flow and favor metastatic extravasation (Fig 3E-F). Quantification performed over a large number of embryos show that sunitinib significantly impairs endothelial remodeling in a dose-dependent manner. Indeed, VEGFR inhibition increases the ratio of intravascular cells and decreases the number of pocketing events (Fig 3E-F). Taken together, these in vivo experiments demonstrate short inhibition of VEGFR is capable of impairing local endothelial remodeling upon CTC arrest without major deleterious effects on hemodynamics and vascular architecture. Thus, VEGFR signaling pathway plays a significant role in the early steps of endothelial remodeling which is required to further drive extravasation of tumor cells.

**Fig. 3.**
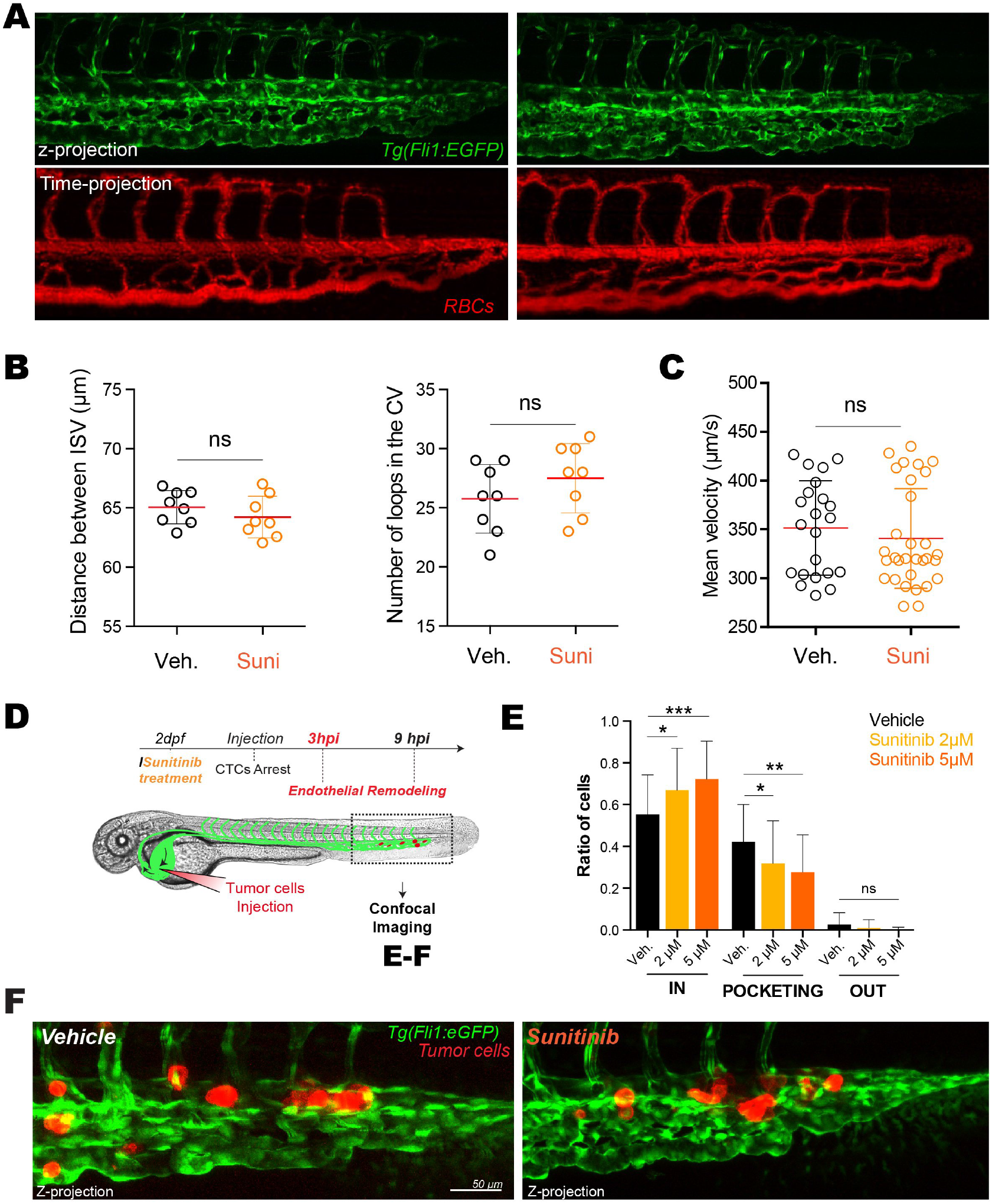
Inhibition of VEGFRs with sunitinib reduces endothelial remodeling in zebrafish embryos. A – Representative confocal image of the caudal plexus architecture (upper panel) and blood flow (lower panel, RBC = red blood cells) 3 hours post-treatment with sunitinib vs vehicle in Tg(fli1a:EGFP; gata1:RFP). B – Quantification of the effect of sunitinib on vessel architecture after 3h of treatment with vehicle or 5μM of sunitinib, N=8, Student t test. C – Blood flow perfusion profiles of the same embryos (from D) and quantification of the blood flow velocity in the caudal plexus of embryos in both conditions after 3h of treatment with vehicle or 5μM of sunitinib. N ZF: vehicle = 8, sunitinib 5μM = 10. D – Experimental setup: zebrafish injection 3hpi (hours post-injection). E – Quantification of intravascular, remodeling and extravasated cells 3hpi. N cells: vehicle = 622, sunitinib 2μM = 375, sunitinib 5μM = 254. N ZF: vehicle = 62, sunitinib 2μM = 34, sunitinib 5μM = 27, Kruskal-Wallis test followed by Dunn’s Multiple Comparison Test. F – Representative image of arrested D2A1 cells in the caudal plexus of vehicle or sunitinib treated embryos.

### Inhibition of the VEGF pathway reduces extravasated of metastatic cells in zebrafish embryos

We next sought to investigate the role of VEGFR signaling during the latter steps of endothelial remodeling (Fig 4A). We thus performed 3D confocal intravital imaging at 9hpi where we previously demonstrated that most of extravasating cells are enwrapped by the endothelium, and thus engaged in the pro-extravasation endothelial remodeling process, or extravasated (Follain et al., 2018a). Inhibition of VEGFRs significantly impairs extravasation of tumor cells at 9hpi (Fig 4B-D). More specifically, while a majority of the tumor cells remained intravascular or in the process of being pocketed upon VEGFR inhibition, they were either fully pocketed or extravasated in control embryos (Fig 4B). To confirm these quantitative results and further investigate endothelial remodeling at high-resolution, we set out a correlative light and electron microscopy (CLEM) experiment on representative embryos in both conditions (Fig 4C-D). CLEM analysis of embryos treated with vehicle immediately after cells injection into the vasculature allowed us to highlight cells that are either in the process of extravasation, through endothelial remodeling, or already fully extravasated and seemingly in contact with the perivascular environment (Fig 4C). Cells undergoing active endothelial remodeling were fully enwrapped with a layer of endothelium observed all around the extravasating CTC (Fig 4Ca). We also observed that extravasated cells were present in the perivascular niche and still in close contact with endothelial cells (Fig 4Cb). When tracking tumor cells in sunitinib-treated embryos, we observed that they remained intravascular with no obvious ultrastructural protrusions that would suggest endothelial remodeling initiation (Fig 4D). Although these cells were in close contact with the endothelium, we could not observe any protrusions from endothelial cells in close proximity suggesting that endothelial remodeling was either impaired or delayed endothelial migration around the arrested CTCs (Fig 4D). We next investigated the effect of sunitinib treatment on the efficiency of extravasation and the formation of micrometastases. We previously demonstrated that the majority of D2A1 cells were extravasated at 24hpi (Follain et al., 2018a). We thus quantified the ratio of extravasated of CTC at 24hpi in embryos that had been treated with vehicle or sunitinib over the whole course of the experiment. In these experiments, we ruled out any deleterious effects on the vascular architecture that could result from high-dose sunitinib and employed sunitinib at low dose (2μM) where only mild perturbations to vascular anatomy could be observed. We did not measure any significant difference in blood flow speed (Fig S2A). Interestingly, in these conditions, the number of extravascular cancer cells was significantly decreased in the caudal plexus region upon VEGFR inhibition (Fig 4E-F, S2B). In details, we measured a 2-fold decrease in extravasation efficiency in sunitinib-treated embryos (Fig 4G) with more CTC staying intravascular compared to vehicle-treated embryos (Fig 4F). Taken together, these in vivo experiments in a validated experimental metastasis assay demonstrate that blood flow indoctrinates endothelial cells through the VEGFR signaling pathway to favor extravasation of tumor cells in zebrafish embryos.

**Fig. 4.**
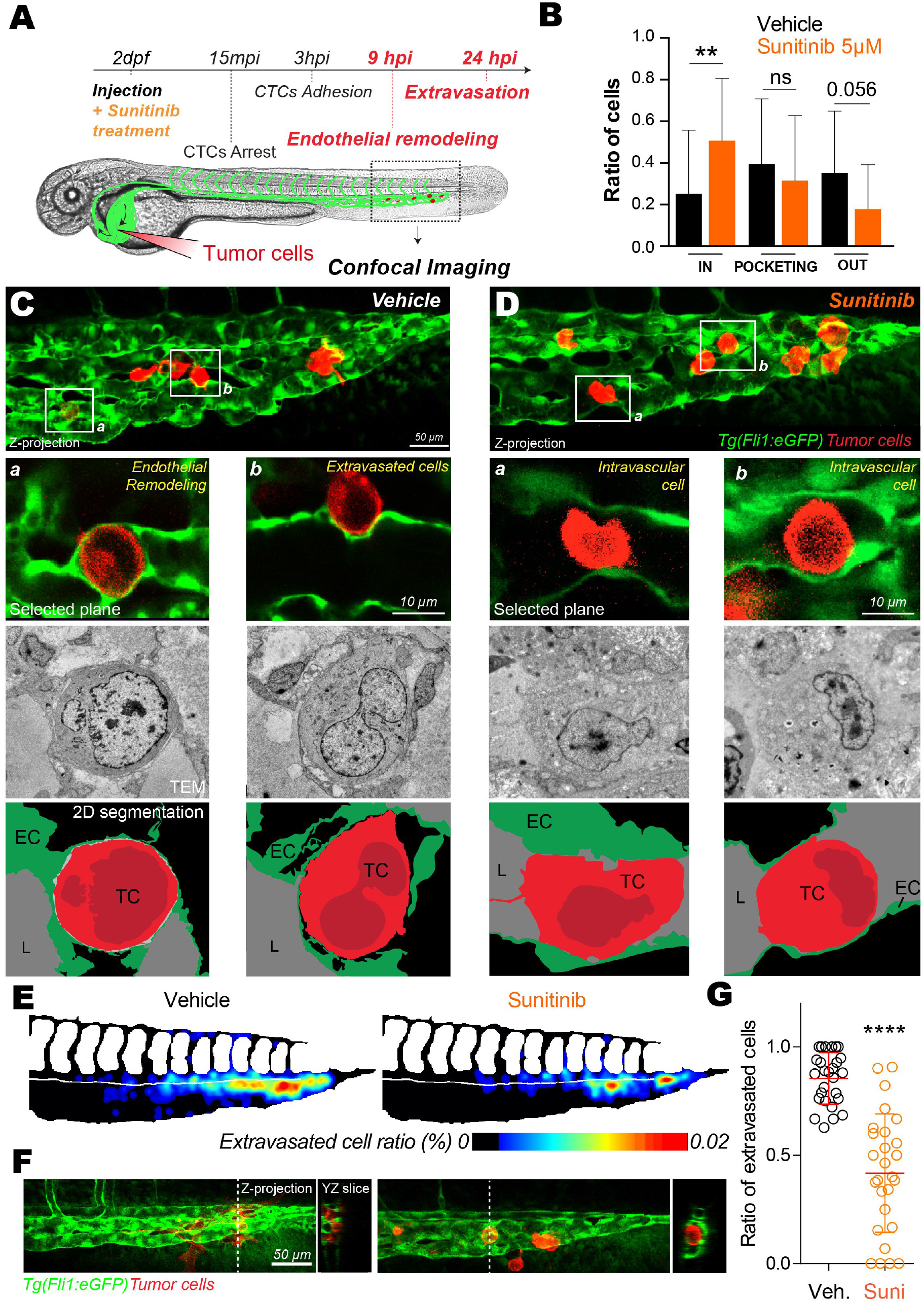
Inhibition of VEGFRs with sunitinib impacts extravasation by endothelial remodeling. A – Experimental setup: zebrafish injection 9hpi. B –Quantification of intravascular, remodeling and extravasated cells 9 hpi. N cells: vehicle = 134, sunitinib = 155. N ZF: vehicle = 23, sunitinib = 25, Kruskal-Wallis test followed by Dunn’s Multiple Comparison Test. C – Representative image of the caudal plexus and associated correlative light and electron microscopy results of tumor cell a and b (white squares on C) in vehicle condition. D - Representative image of the caudal plexus and associated correlative light and electron microscopy results of tumor cell c and d (white squares on D) in sunitinib condition. E – The heatmaps show the quantification and location of extravascular CTCs at 24 hpi in the caudal plexus treated with vehicle or 2μM of sunitinib. F – Representative images and orthoslice at 24hpi. G – Quantification of extravasated cells ratio at 24 hpi treated with vehicle or 2μM of sunitinib. N cells: vehicle = 588, sunitinib = 441. N ZF: vehicle = 27, sunitinib = 27, Mann Whitney test.

## Discussion

CTC extravasation is an essential step preceding the rate-limiting organ colonization and the foundation of secondary tumor foci (Valastyan and Weinberg, 2011). Yet, the cellular dynamics and molecular mechanisms of CTC extravasation remain incompletely understood. Extravasation of tumor cells is a rare event at the scale of an entire organism and full understanding requires resolutive imaging technologies applied to relevant experimental metastasis models. Based on this, we previously discovered that extravasation occurs mainly through an active endothelial-dependent process that we named endothelial remodeling (Follain et al., 2018a). Our findings were recently extended to human melanoma and cervix cancer lines (Allen et al., 2019), suggesting that it is not a cell line specific mechanism. Incidentally, a similar process had also been linked to extravasation of stem cells (Allen et al., 2017) and is at play during the important step of blood clot removal, through angiophagy (Lam et al., 2010; Grutzendler et al., 2014). While recent work demonstrated that endothelial cell death through necroptosis is important for tumor cell extravasation (Strilic et al., 2016), earlier work had also eluded to endothelialization of tumor cells during metastasis (Lapis et al., 1988; Paku et al., 2000). Altogether, this demonstrates that the endothelial wall is a key determinant of the metastatic success. Yet, the underlying molecular triggers and mechanisms favoring plasticity of the endothelium during extravasation remain poorly understood. Using a flow-tuning approach in the zebrafish embryo and in vitro microfluidics, we demonstrated that endothelial remodeling is driven and tuned by hemodynamic cues from the blood flow (Follain et al., 2018a). Here, using a combination of in vitro microfluidics and transcriptomics, but also of intravital imaging and ultrastructural analysis in the zebrafish embryo, we demonstrate that endothelial remodeling around arrested tumor cell is driven, in part, by VEGFR signaling downstream of hemodynamic cues. This is further supported by the observation that endothelial remodeling can be impaired by sunitinib (also known as Sutent©), a potent inhibitor of VEGFR pathway used to prevent neo-angiogenesis during cancer treatment. We thus demonstrate that intravascular cues that are at play during metastasis are capable of indoctrinating the endothelial wall through a classical VEGFR pathway to stimulate metastatic extravasation of CTCs. Using transcriptomic analysis, we first demonstrate that permissive flow profiles favor the activation of molecular programs that are linked to angiogenesis, as previously described during developmental angiogenesis (García-Cardeña et al., 2001; Galie et al., 2014; Nakajima et al., 2017; Nicoli et al., 2010). Within this molecular program, VEGF receptors appear significantly upregulated in response to flow. Endothelial cells have developed several strategies to sense flow which include the cilium, the glycocalyx, mechano-sensitive ions channels, G protein coupled receptors, caveolae, adhesion receptors and VEGFRs (Freund et al., 2012; Baratchi et al., 2017). Thus, mechanisms that are relevant for developmental angiogenesis are also likely to be essential for pathological vessel remodeling, including during metastatic extravasation. With the idea to further validate a molecular node that controls extravasation of tumor cells through endothelial remodeling, we decided to employ sunitinib, as a potent endothelial remodeling inhibitor. Sunitinib treatment successfully inhibits the extravasation of tumor cells both in an in vitro microfluidic setup and in the zebrafish embryos. Our data argue that the VEGFR pathway is a major actor of the flow-dependent endothelial remodeling, driving extravasation as we described previously (Follain et al., 2018a). Whether VEGFR, and more specifically the mechanosensory VEGFR2 and 3, are direct flow mechanotransducers, are acting together with other mechanosensory complexes or are activated downstream of other flow sensors remains to be elucidated (Freund et al., 2012). The VEGFR canonical activation is complex, due to partial ligand specificity and crosstalk between VEGF A, B, C, D or PIGF ligands and VEGFR1, R2 or R3 receptors and the heterodimerization between the VEGFRs (Domigan et al., 2015; Simons et al., 2016). Interestingly, non-canonical (ligand-independent) VEGFR activation has been described in the context of mechano-transduction (Tzima et al., 2005; Conway et al., 2013; Baeyens et al., 2015; Coon et al., 2015). Blood flow changes that occur at the apical side of the endothelial cell, when CTC arrest, might be sufficient to activate VEGFR kinase cascade and transduce the signal into the endothelial cells (dela Paz et al., 2012; Koch et al., 2011). Alternatively, indirect forces that result from shear forces on the arrested CTC and that could be transmitted to the endothelial cell which is directly engaged through CTC-endothelium cell-cell adhesions on the luminal side are likely to favor certain transcriptional programs (Schaefer and Hordijk, 2015; Baeyens and Schwartz, 2016). Such a hypothesis is exciting and further work is needed to determine whether indirect flow forces on arrested tumor cells are likely to impact neighboring endothelial cells. Combination of biophysical tools (such as optical tweezing technologies to manipulate CTC and/or endothelial cells) with microfluidic approaches could be helpful in that regard (Follain et al., 2018a; Harlepp et al., 2017; Osmani et al., 2019). In addition, there are obviously additional molecular programs that are likely to be involved during this endothelium-dependent process. Interestingly, the process of angiophagy is driven by MMP2/9 activity (Lam et al., 2010) and could be involved in driving endothelial remodeling that highly depends on endothelial cell protrusions. In our case, however, expression levels of MMP 2 and 9 are only mildly affected (data not shown). Furthermore, more work is needed to demonstrate that whether endothelial remodeling is a universal process used by CTCs to extravasate, or whether this is limited to certain vascular and hemodynamic conditions. The latter likely depend on the organ of metastasis. Although the vasculature of the early zebrafish embryo differs from mature vessels of other relevant models such as mice, we had shown that endothelial remodeling is also at play during metastatic extravasation of tumor cells in the mouse brain. In that case, VEGFR or other signaling pathways could control endothelial remodeling and metastatic extravasation. More work is required to determine whether endothelial remodeling occurs in distinct vascular environments and whether they use common or alternative signaling pathways. Sunitinib is a tyrosine-kinase receptor inhibitor, mainly targeting VEGFR, PDGFR and c-KIT receptors that was first identified in 1999 (Fong et al., 1999). It is currently clinically used for several cancer treatments including renal or gastrointestinal cancer (Younus et al., 2010). Several clinical trials are being conducted to block vascular development around well-established tumors and in comparison with similar drugs such as Pazopanib, another VEGFR/PDFGR/cKit Inhibitor (Rodriguez-Vida et al., 2017; Chellappan et al., 2017). Despite positive results in its ability to impair tumor growth and cancer progression (Young et al., 2010), the clinical impact of sunitinib is currently questioned as contradictory results have been obtained in the context of tumor metastasis. At this stage, although sunitinib displays detrimental properties that impair its usage in this context, we believe that its powerful inhibition of the VEGFR pathway demonstrates that metastatic extravasation of tumor cells is likely to occur through VEGFR-dependent endothelial remodeling. In line with our results, a recent study used zebrafish model to predict the clinical efficiency of bevacizumab, another anti-angiogenic therapeutic agent, in antimetastatic response (Rebelo de Almeida et al., 2020). Inhibition of Vascular endothelial growth factor-A (VEGF-A) also induced long-term dormancy of lung cancer micrometastases by preventing angiogenic growth to macrometastases in a mouse model of brain metastasis (Kienast et al., 2010). Flow-independent biomechanical cues might also be elemental to the endothelial response during metastatic progression. It was recently demonstrated that stiffness reduction in liver metastasis induced a higher therapeutic response to bevacizumab (Shen et al., 2020), similarly to our observations pointing at a crosstalk between VEGFR-mediated endothelial response and external biomechanical cues. More work is needed to identify additional signaling pathways, and more specific molecules, to efficiently counteract this new mechanism of metastatic extravasation. Altogether, our data argue that the endothelial remodeling leading to extravasation of metastatic cells can be chemically inhibited in vivo. We are confident that molecular targeting of this step (i.e endothelial remodeling) is at reach and that it will provide new therapeutic means to prevent, efficiently, metastatic progression at the extravasation step.

## Acknowledgements

We thank all members of the Goetz Lab for helpful discussions and Sebastien Harlepp for assistance throughout the study. We are very grateful to Francesca PERI and Kerstin RICHTER (EMBL) for providing zebrafish embryos at the early stage of the project. We thank the imaging facility members from EFS Strasbourg, IGBMC and INCI for access to light and electron microscopes.

## Author Contributions

GF and NO performed most of the experiments and analysis with help from LM and MJGL. OL performed RT-qPCR analysis and generated stable cell lines. IB performed the CLEM procedures with input from GF and LM. AP, NP, RC and SB performed the RNAseq experiment and sequence alignment. SH supervised the study and performed imaging and blood flow analysis. GF and NO wrote the manuscript with input from OL and JGG. JGG conceived the project and supervised the study.

## Material Methods

### Cell lines

D2A1 were provided by Robert A. Weinberg (MIT). Cells stably expressing LifeAct-TdTomato or Luciferin were obtained using lentiviral transfection. Cells were grown as previously described (Shibue et al., 2012), in DMEM with 4.5 g/l glucose (Dutscher) supplemented with 5per cent FBS, 5percent NBCS, 1percent NEAA and 1percent penicillinstreptomycin (Gibco). Human Umbilical Vein Endothelial Cells (HUVEC) (PromoCell) were grown in ECGM (PromoCell) supplemented with supplemental mix (PromoCell C-39215) and 1percent penicillin-streptomycin (Gibco). To maximize the reproducibility of our experiments, we always used these cells at 4th passage in the microfluidic channels.

### Antibodies

mouse anti-human CD31(PECAM) monoclonal antibody (MEM-5) was purchased from Thermo, mouse anti-human FLT1 (ab9540) and rat anti-mouse FLT4 (ab51874) monoclonal antibodies were purchased from AbCam, rabbit anti-KDR monoclonal antibody (D5B1) was purchased from Cell Signaling.

### Zebrafish handling and sunitinib treatment

Tg(fli1a:EGFP), Tg(fli1a:EGFP; gata1:RFP) and Tg(CD41:EGFP; gata1:RFP) zebrafish (Danio rerio) embryos were used in the experiments. Embryos were maintained in Danieau 0.3X medium (17,4 mM NaCl, 0,2 mM KCl, 0,1 mM MgSO4, 0,2 mM Ca(NO3)2) buffered with HEPES 0,15 mM (pH = 7.6), supplemented with 200 μM of 1-Phenyl-2-thiourea (Sigma-Aldrich) to inhibit the melanogenesis, as previously described (Goetz et al., 2014). Sunitinib (Sigma-Aldrich) was added in the breeding water of the embryos directly after injection of tumor cells at the given concentration of 2 μM or 5 μM vs vehicle (DMSO).

### Control experiments for sunitinib effect on zebrafish embryos

Sunitinib treatment was tested for its potential impact on vascular architecture and/or on blood flow pattern in the caudal region. After 3 or 24 hours of treatment (sunitinib or vehicle), we performed confocal microscopy using an inverted TCS SP5 confocal microscope with a 20x/0,75 (Leica) and manually analyzed the architectures of landmark vessels. In the plexus, the conservation of the distance between intersegmental vessels and the number of vascular branching in the caudal veins was quantified. Also, we performed fast recording using the resonant scanner at a speed of 27 fps in Tg(CD41:EGFP; gata1:RFP) embryos. We measured the mean flow velocity using the TrackMate plugin in Fiji (Tinevez et al., 2017).

### Intravascular injection and imaging of CTCs in the zebrafish embryo

48 hour post-fertilization (hpf) Tg(Fli1a:EGFP) embryos were mounted in 0.8percent low melting point agarose pad containing 650 μM of tricain (ethyl-3-aminobenzoate-methanesulfonate) to immobilize the embryos. D2A1 LifeAct-TdTomato cells were injected with a Nanoject microinjector 2 (Drummond) and microforged glass capillaries (25 to 30 μm inner diameter) filled with mineral oil (Sigma). 18nL of a cell suspension at 100.106 cells per ml were injected in the duct of Cuvier of the embryos under the M205 FA stereomicroscope (Leica). For caudal plexus, confocal imaging was performed with an inverted TCS SP5 confocal microscope with a 20x/0,75 (Leica). The caudal plexus region (around 50μm width) was imaged with a z-step of less than 1.5 μm for at least 20 embryos per conditions from 3 independent experiments. For zebrafish head imaging, a multiphoton microscope Bruker Ultima Investigator equipped with a 25x/1.10 water immersion (Nikon) and an Insight X3 pulsed laser (Spectra Physics) was used. Cell number and situations was manually characterized (Intravascular, ongoing endothelial remodeling/pocketing, extravascular) using z-projections and orthogonal views in ImageJ. Correlative Light and Electron Microscopy was performed to describe ultrastructural characteristics of CTCs and the endothelium in the zebrafish embryo. Chosen embryos of both condition (Vehicle and Sunitinib treated) were imaged using confocal microscopy between 3 to 4 hpi. Just after imaging, they were chemically fixed and processed for EM (see dedicated section “EM preparation”).

### EM preparation and Correlation between light and electron microscopy

The samples have been post fixed in a solution of 2,5percent glutaraldehyde and 2percent paraformaldehyde in 0.1 M Cacodylate buffer at 4°C overnight. Samples were rinsed in 0.1M Cacodylate buffer for 2×5min and post-fixed using 1 per cent OsO4 in 0.1M Cacodylate buffer, for 1h at room temperature. Then, samples were rinsed for 3×5min in 0.1M Cacodylate buffer. Followed by washing 2×5min in pure water. Then, secondary post-fixed with 4 per cent water solution of uranyl acetate, 1h at room temperature. Followed by 5 min wash in pure water, the samples were stepwise dehydrated in Ethanol (50 per cent 3×5 min, 70 per cent 3×10, 90 per cent 3×10 and 100 per cent 3×10) and infiltrated in a graded series of Epon (Ethanol abs/Epon 3/1, 1/1, 1/3, each 30min). Samples were left in absolute Epon (EmBed812 - EMS) overnight. Then, samples were placed in a fresh absolute Epon for 1h and polymerized (flat embedded) at 60°C for 24-48h. Once polymerized, most surrounding Epon was cut off using razorblade and samples were mounted on empty Epon blocks (samples flat at the surface of the blocks) and left at 60 °C for 24h-48h. To do the correlation in 3d, semi thin sections (500nm) were obtained using glass knife in ultramicrotome LEICA UCT. Sections were placed on slide, stained with 1 per cent borax Toluidine blue solution and checked out in the optical microscope. For zebrafish embryos, we applied the same protocol and used anatomical landmark to retrieve the ROI (Dorsal aorta, tumor cells…). Ultrathin sections (100nm) were serially sectioned using ultramicrotome (Leica Ultracut UCT), collected on formvar-coated slot grids and stained with 4 per cent water solution of uranyl acetate for 5min and lead citrate for 3min. Ultra-thin sections were imaged with a CM120 transmission electron microscope (Philips Biotwin) operating at 120 kV. Images were recorded with Veleta 2k x 2k (Olympus-SIS) camera using iTEM software.embedded and the ROI approached by serial sectioning.

### Microfluidic experiments

For endothelial remodeling experiments in vitro, two μ-slides I0.4 Luer (IBIDI) coated with fibronectin from bovine plasma at 10μg/ml (Sigma F-1141) were used in parallel for each experiment. HUVEC cells were seeded at 80 000 cells per channel. Medium was changed twice a day for 2 or 3 days, before perfusing the channels under a flow of 400μm/sec using REGLO Digital MS-2/12 peristaltic pump (Ismatec) and Tygon tubbing (IDEX). Sunitinib treatment was added at a concentration of 0,1μM in flow. At confluence, D2A1 LifeAct-TdTomato cells were added at a concentration of 200 000 cells/ml for 10 min. Then, floating tumor cells were washed using fresh medium and the channels were incubated for 16 h with flow. Position of the tumor cells and presence of endothelial remodeling around tumor cells relative to the HUVEC monolayer were determined using the piezzo stage of the confocal microscope after fixation and Immunofluorescent staining (see next section).

### Immunofluorescent staining in the microfluidic channels

Cells were fixed using 4 per cent PFA (Electronic Microscopy Sciences), permeabilized with 0.2 per cent Triton-X100 (Sigma) and quenched with 2mg/ml NaBH4 (Sigma) 10 min at room temperature before using the following primary antibodies. Following secondary antibodies were used: goat anti-mouse/rat/rabbit coupled with Alexa Fluor 488, Cy3, Alexa 555, Cy5 or Alexa647 (Invitrogen). Cells were mounted using Vectashield (Vector Laboratories).

### RNA sequencing

3 independent couples (flow and no flow) of HUVEC samples were isolated from IBIDI μ-slides I0,4 Luer (IBIDI) using Tri-reagent (MRC) 100 μl of Tri-reagent was added directly in one side of the channel and aspire in the other side 5 times. This was followed by chloroform extraction and alcohol washing. Total cDNA was obtained using Thermo Fisher kit (SuperScriptTM VILOTM Master mix). RNA integrity was assessed with the Agilent total RNA Pico Kit on a 2100 Bioanalyzer instrument (Agilent Technologies, Paolo Alto, USA). The construction of libraries was done with the “SMARTer® Stranded Total RNA-Seq Kit v2 - Pico Input Mammalian” (TaKaRa Bio USA, Inc., Mountain View, CA, USA) with a final multiplexing of 12 libraries according to the manufacturer’s instructions. The library pool was denatured according to the Illumina protocol “Denature and Dilute Libraries Guide” and then deposited at a concentration of 1.3 pM to be sequenced on the NextSeq 500 (Illumina Inc., San Diego, CA, USA). The transcriptome data set, composed of sequencing reads, was generated by an Illumina NextSeq instrument. The objective is to identify genes that are differentially expressed between two experimental conditions: flow and no flow. First, the data were mapped to the human genome/transcriptome (hg19) using the hisat2 software (Kim et al., 2015), a fast and sensitive alignment program. The total reads mapped were finally available in BAM format for raw read counts extraction. Next, read counts were found by the htseq-count tool of the Python package HTSeq (Anders et al., 2015) with default parameters to generate an abundant matrix. Then, differential analyses were performed by the DESEQ2 (Love et al., 2014) package of the Bioconductor framework. Up-regulated and down-regulated genes were selected based on the adjusted p-value cutoff 10 per cent. Finally, Gene Ontology Consortium (http://www.geneontology.org/) platform was used for data analysis and heatmaps creation. The heatmaps were formatted using IGOR software.

### qPCR validation

3 to 7 independent couples (flow and no flow) of HUVEC samples total RNA were isolated from IBIDI μ-slides I0,4 (Luer Family) using Tri-reagent (MRC) followed by chloroform extraction and alcohol precipitation. Total cDNA was obtained using ThermoFisher kit (SuperScriptTM VILOTM Master mix). RT qPCR reactions were made using either TaqMan master mix (ThermoFisher - 4444557) or SYBR green master mix (ThermoFisher – A25742) in Applied Biosystem qPCR machine (QuantStudioTM 3 - ThermoFisher). See next table of qPCR primer sequences for Flt1, Kdr, Flt4, Notch1, Lama4 (lab designed). For Gapdh, a commercial TaqMan probe was used (ThermoFisher - 4333764F). Amplification results were normalized using Gapdh level and double ΔcT method. Targets Primer sequences

h Flt1 fwd CCA GCA GCG AAA GCT TTG CG
h Flt1 rev CTC CTT GTA GAA ACC GTC AG
h Kdr fwd ATG ACA TTT TGA TCA TGG AGC
h Kdr rev CCC AGA TGC CGT GCA TGA G
h Flt4 fwr TGC AAG AGG AAG AGG AGG TCT
h Flt4 rev CAG GCT TGG CGG GCT GTC C
h Notch1 fwd CAG GAC GGC TGC GGC TCC TAC
h Notch1 rev CCG CCG TTC TTG CAG GGC GAG
h Lama4 fwr ACC TCC TCA ATC AAG CCA GA
h Lama4 rev TCA GCC ACT GCT TCA TCA CT

### Western blotting

Extracts corresponding to similar cell numbers were loaded on 4-20 per cent polyacrylamide gels (Biorad) and run under denaturing conditions. The following primary antibodies were used: FLT1 (ab9540 - abcam), KDR (D5B1 – Cell Signaling), FLT4 (ab51874 - abcam), α-Tubulin (DM1A - Calbiochem). HRP-conjugated secondary antibodies were used with ECL (GE Healthcare) for reading using the ChemiDoc XRS (Biorad). Intensities were normalized over α-Tubulin levels.

### Stastistical analysis

Statistical analysis was performed using the GraphPad Prism program version 5.04. The Shapiro-Wilk normality test was used to confirm the normality of the data. For data following a Gaussian distribution, a Student unpaired two-tailed t test, with Welch’s correction in case of unequal variances was used. For data not following a Gaussian distribution, the Mann-Whitney test were used. When more than 3 groups are compared, a Kruskal-Wallis test followed by Dunn’s Multiple Comparison Test was used. For qualitative data, the Fisher test was used. Illustrations of these statistical analyses are displayed as the mean +/− standard deviation (SD). p-values smaller than 0.05 were considered as significant. *, p<0.05, **, p<0.01, ***, p<0.001, ****, p<0.0001, ns=not significant

---

**Fig. 5. Supplemental Figure 1:**
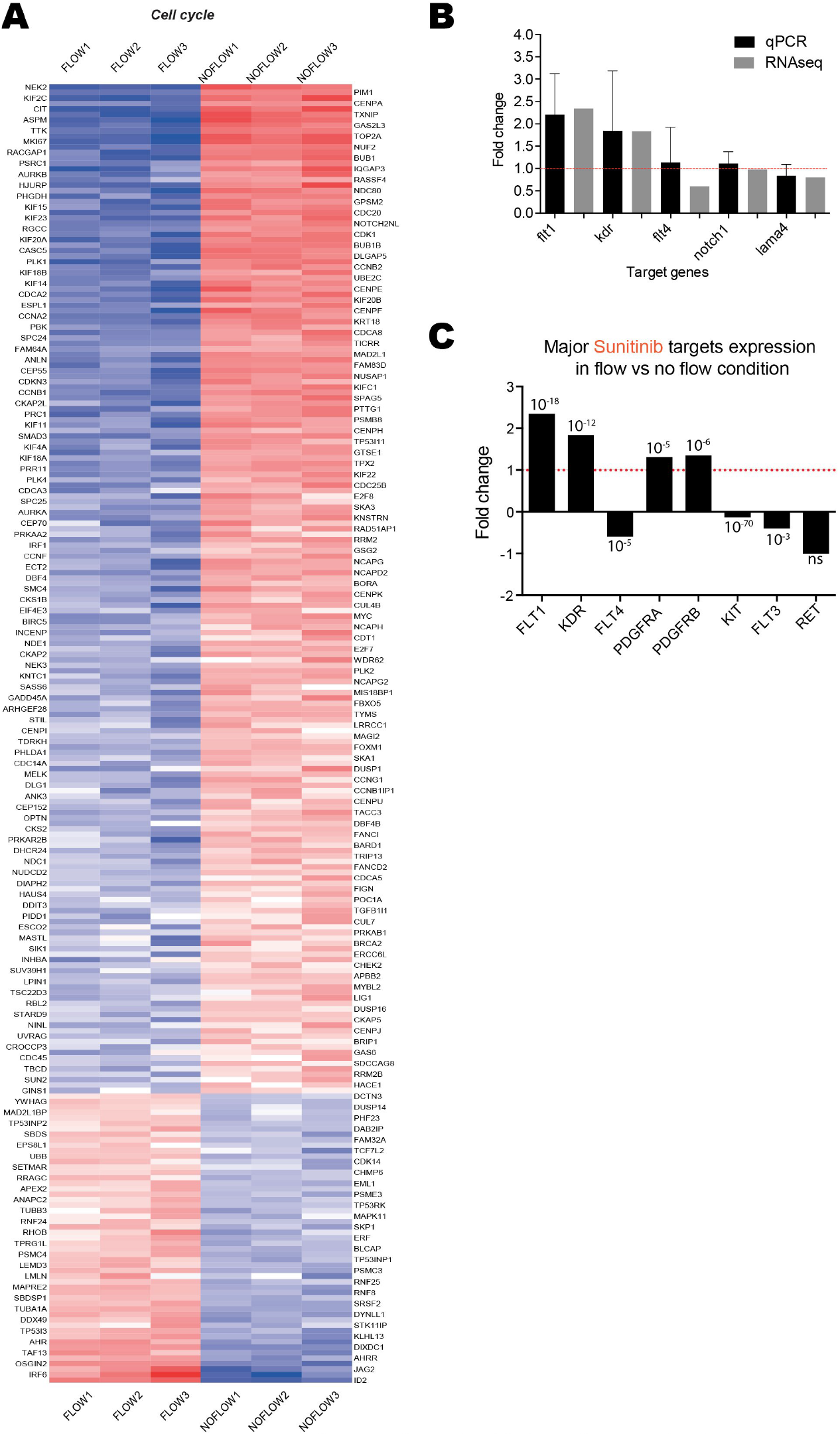
Gene expression in HUVEC cells treated with or without flow. A – Fold changes heatmap, based on GO class: Cell Cycle, showing significantly impacted genes in flow condition compared to no flow. B – RT-qPCR data confirming RNA sequencing results. Graphic shows the comparison between RT-qPCR and RNAseq. C – Fold change of expression level (RNAseq) of the major sunitinib targets in flow vs no flow condition (numbers are adjusted p-values)

**Fig. 6. Figure S2:**
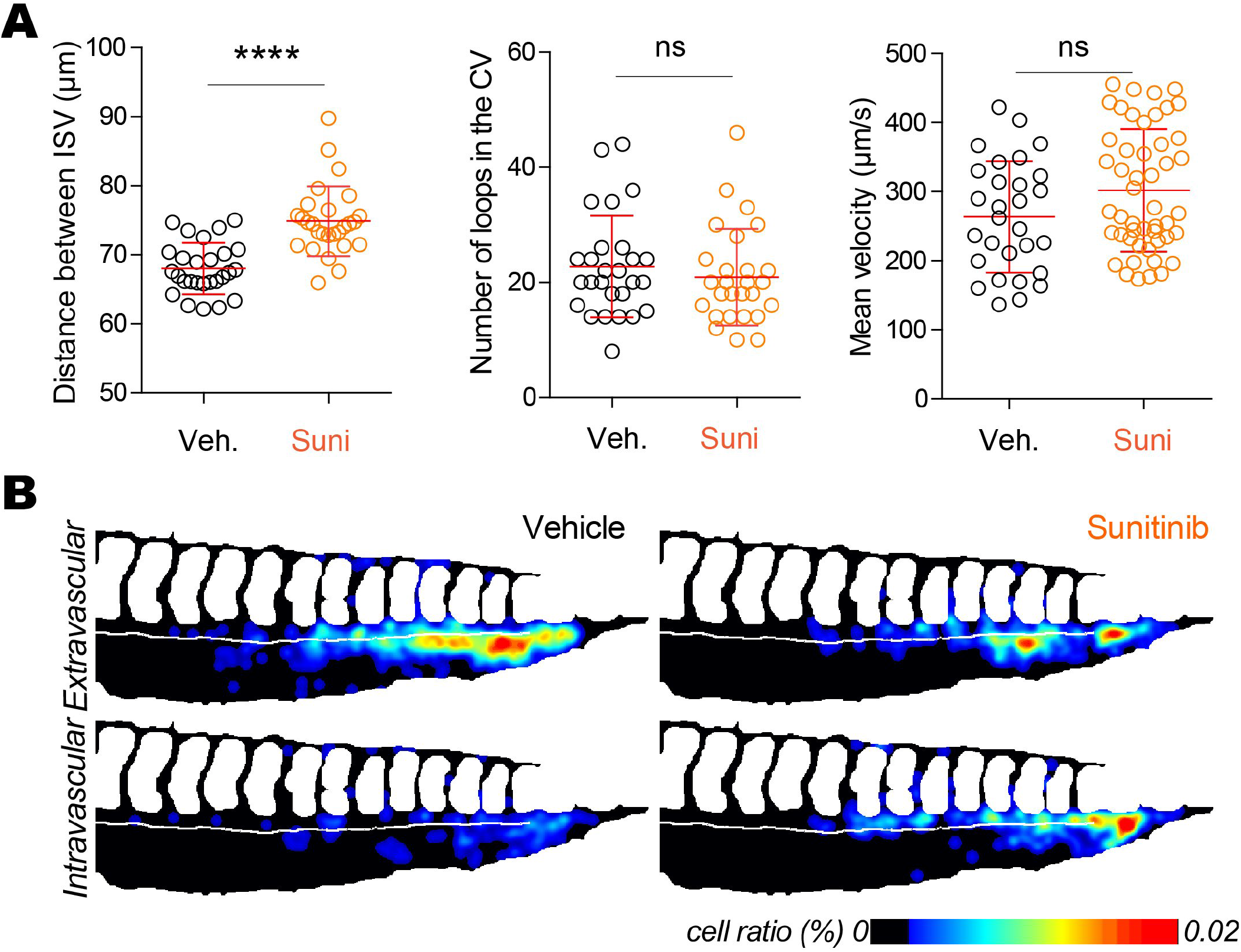
Inhibition of VEGFRs with sunitinib reduces extravasation in the caudal plexus. A – Quantification of the effect of sunitinib on vessel architecture after 24h of treatment, N ZF: vehicle = 27, sunitinib = 27, Mann Whitney test. B – The heatmaps show the quantification and location of CTCs at 24 hpi in the caudal plexus. Quantification of the blood flow velocity in the caudal plexus of embryos in both conditions after 24h of treatment with vehicle or 2μM of sunitinib. N ZF: vehicle = 13, sunitinib 2μM = 18.

